# Exercise Training and Cold Exposure Trigger Distinct Molecular Adaptations to Inguinal White Adipose Tissue

**DOI:** 10.1101/2023.10.16.562635

**Authors:** Maria Vamvini, Pasquale Nigro, Tiziana Caputo, Kristin I. Stanford, Michael F. Hirshman, Roeland J.W. Middelbeek, Laurie J. Goodyear

**Affiliations:** Section on Integrative Physiology and Metabolism, Joslin Diabetes Center, Harvard Medical School, Boston, MA; Division of Endocrinology, Diabetes and Metabolism, Beth Israel Deaconess Medical Center, Harvard Medical School, Boston, MA; Department of Physiology and Cell Biology, Diabetes and Metabolism Research Center, Dorothy M. Davis Heart and Lung Research Institute, The Ohio State University Wexner Medical Center, Columbus, OH, USA

**Keywords:** exercise, cold, adipose tissue, transplantation, proteomics, secretome

## Abstract

Exercise training and cold exposure both improve systemic metabolism, but the mechanisms are not well-established. We tested the hypothesis that adaptations to inguinal white adipose tissue (iWAT) are critical for these beneficial effects by determining the impact of exercise-trained and cold-exposed iWAT on systemic glucose metabolism and the iWAT proteome and secretome. Transplanting trained iWAT into sedentary mice improved glucose tolerance, while cold-exposed iWAT transplantation showed no such benefit. Compared to training, cold led to more pronounced alterations in the iWAT proteome and secretome, downregulating >2,000 proteins but also boosting iWAT’s thermogenic capacity. In contrast, only training increased extracellular space and vesicle transport proteins, and only training upregulated proteins that correlate with favorable fasting glucose, suggesting fundamental changes in trained iWAT that mediate tissue-to-tissue communication. This study defines the unique exercise training- and cold exposure-induced iWAT proteomes, revealing distinct mechanisms for the beneficial effects of these interventions on metabolic health.

**GRAPHICAL ABSTRACT:** 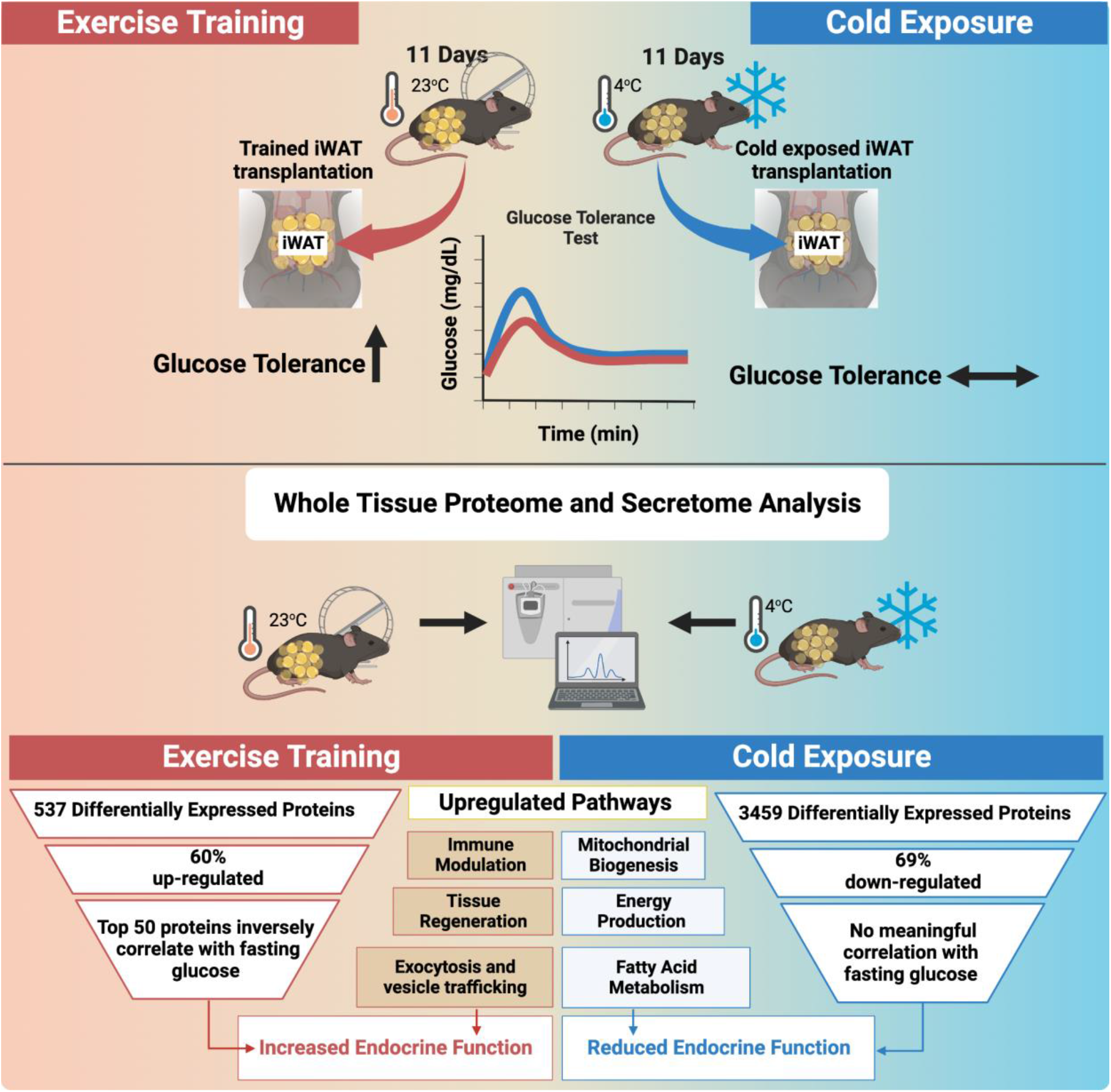

## INTRODUCTION

Exercise training is established as an effective strategy for helping prevent obesity and type 2 diabetes^1^. Regular exercise promotes improvements in systemic metabolism including increased insulin sensitivity, lipid metabolism, and energy expenditure, as well as having beneficial adaptations in most organs and cells in the body ^2^. Recent studies demonstrate that of these organs, exercise training has especially robust effects on subcutaneous white adipose tissue (WAT), dramatically altering gene and protein expression, causing tissue remodeling, and promoting vascularization and innervation ^3–7^; adaptations that lead to a more favorable metabolic profile ^3^.

Our previous work demonstrated that transplanting subcutaneous inguinal WAT (iWAT) from exercise-trained mice into recipient mice leads to a significant improvement in systemic glucose regulation^3^. This is likely due to the transplanted tissue exerting endocrine effects on other tissues, resulting in increased glucose uptake in brown adipose tissue and skeletal muscles^3^. Similar to exercise training, cold exposure elicits profound adaptations in various tissues, including iWAT ^8–11^. Cold exposure stimulates the activation of brown adipose tissue (BAT)^12,13^ and the emergence of beige adipocytes within WAT depots, resulting in enhanced thermogenesis and energy expenditure ^14,15^. Whether iWAT adaptations induced by cold exposure can yield benefits on glucose metabolism similar to exercise training is not known, but is important to investigate because it would help delineate new metabolic pathways, which, in concert with exercise, could be targeted for more precise therapeutic interventions in the management of metabolic disorders such as type 2 diabetes and obesity ^16^.

Here, we compared the effects of transplanting exercise trained versus cold-exposed iWAT into sedentary recipient mice on glucose tolerance. Unexpectedly, we found that glucose tolerance in recipient mice was only improved in mice transplanted with trained-iWAT and not cold-exposed iWAT. To understand the molecular underpinnings of these differences, we employed histological assessment of iWAT as well as proteome analysis of both iWAT and its secretome. We find striking differences between training and cold exposed iWAT, including unique protein and molecular signatures.

## RESULTS

### Transplantation of subcutaneous inguinal white adipose tissue (iWAT) from exercise-trained and cold-exposed mice

To determine whether exercise training and cold exposure have the same beneficial effects on glucose homeostasis, male mice were given free access to an exercise wheel or were housed individually in standard cages for 11 days at room temperature or cold incubators at 4°C. Subcutaneous iWAT (0.85 g) was removed and transplanted into the visceral cavity of 12-week-old sedentary recipient male mice (Figure 1A). Nine days post-transplantation there was a significant improvement in glucose tolerance in mice transplanted with iWAT from exercise-trained mice. In contrast, there was no change in glucose tolerance in mice transplanted with iWAT from sedentary or cold-exposed mice (Figure 1B). These data suggest that there are unique characteristics of the trained iWAT not present in the cold exposed iWAT which may confer beneficial effects on glucose homeostasis.

**Figure 1.**
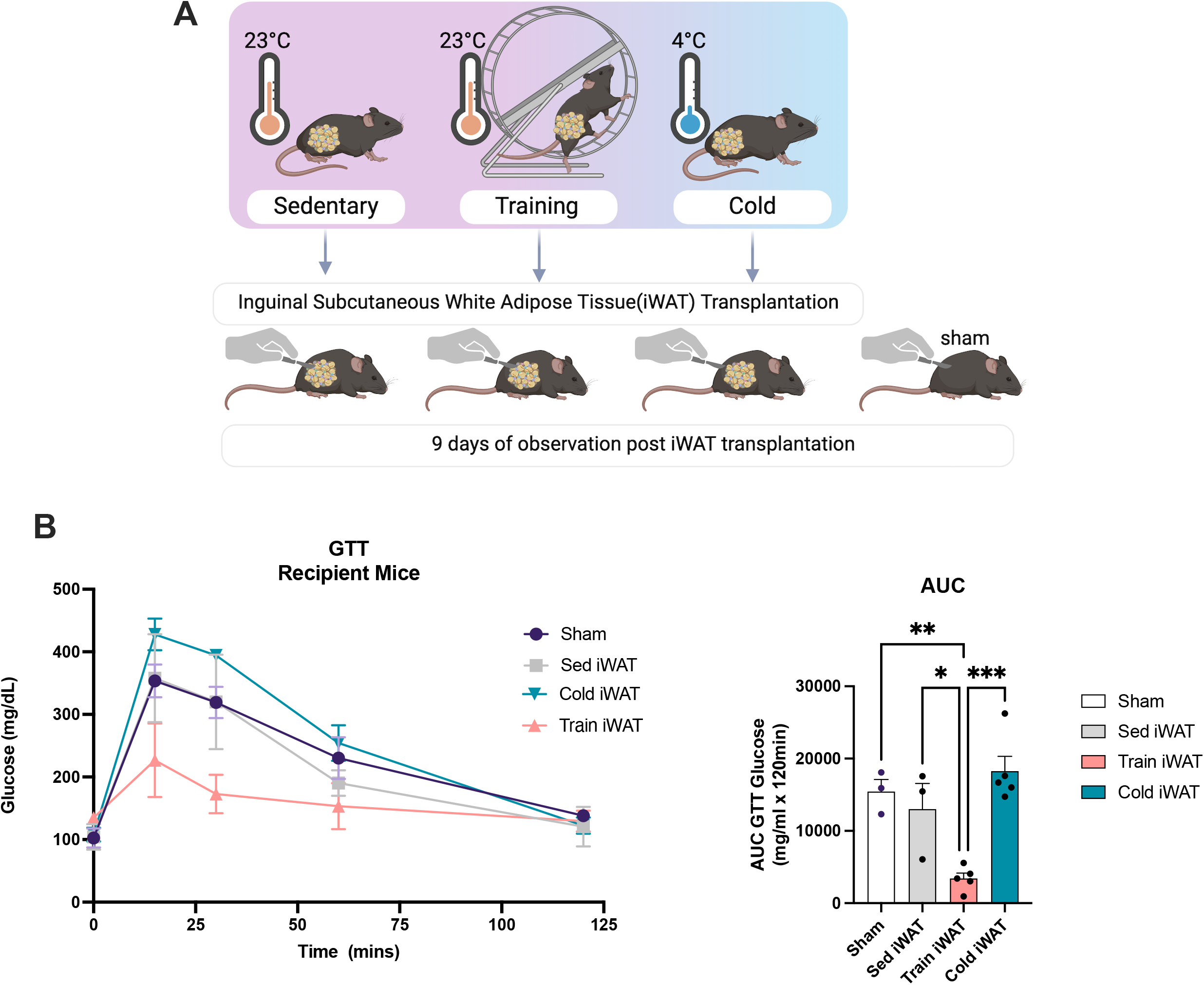
iWAT transplantation from exercise trained but not cold-exposed mice improved glucose tolerance. (A) iWAT was collected from male mice after 11 days of exercise, cold exposure, or a sedentary lifestyle and transplanted into male C57BL6 recipient mice. All analyses were conducted nine days post-transplantation. (B) Intraperitoneal glucose tolerance test (ipGTT) and area under the curve (AUC) in recipient mice after 9 days from transplantation. Data are presented as mean±SEM and were compared using One-way ANOVA. *p<0.05,**p<0.01, and ***p<0.001

### Phenotypic responses of mice and iWATs to exercise and cold

To investigate the effects of exercise training and cold exposure on mouse phenotype and iWAT adaptations, mice underwent the same 11-day exercise training and cold protocols described above for transplantation (Figure 2A). The average running distance for the trained mice was 6.1+/-2.5 Km/day. Sedentary mice exhibited a significantly greater increase in body weight over the 11 days compared to both exercise-trained and cold-exposed mice and there was no difference in body weights between the trained and cold-exposed groups (Figure 2B). Cold exposure resulted in significantly higher food intake compared to both sedentary and trained mice (Figure 2C), whereas only trained mice showed improved fasting glucose concentrations compared to sedentary mice (Figure 2D).

**Figure 2.**
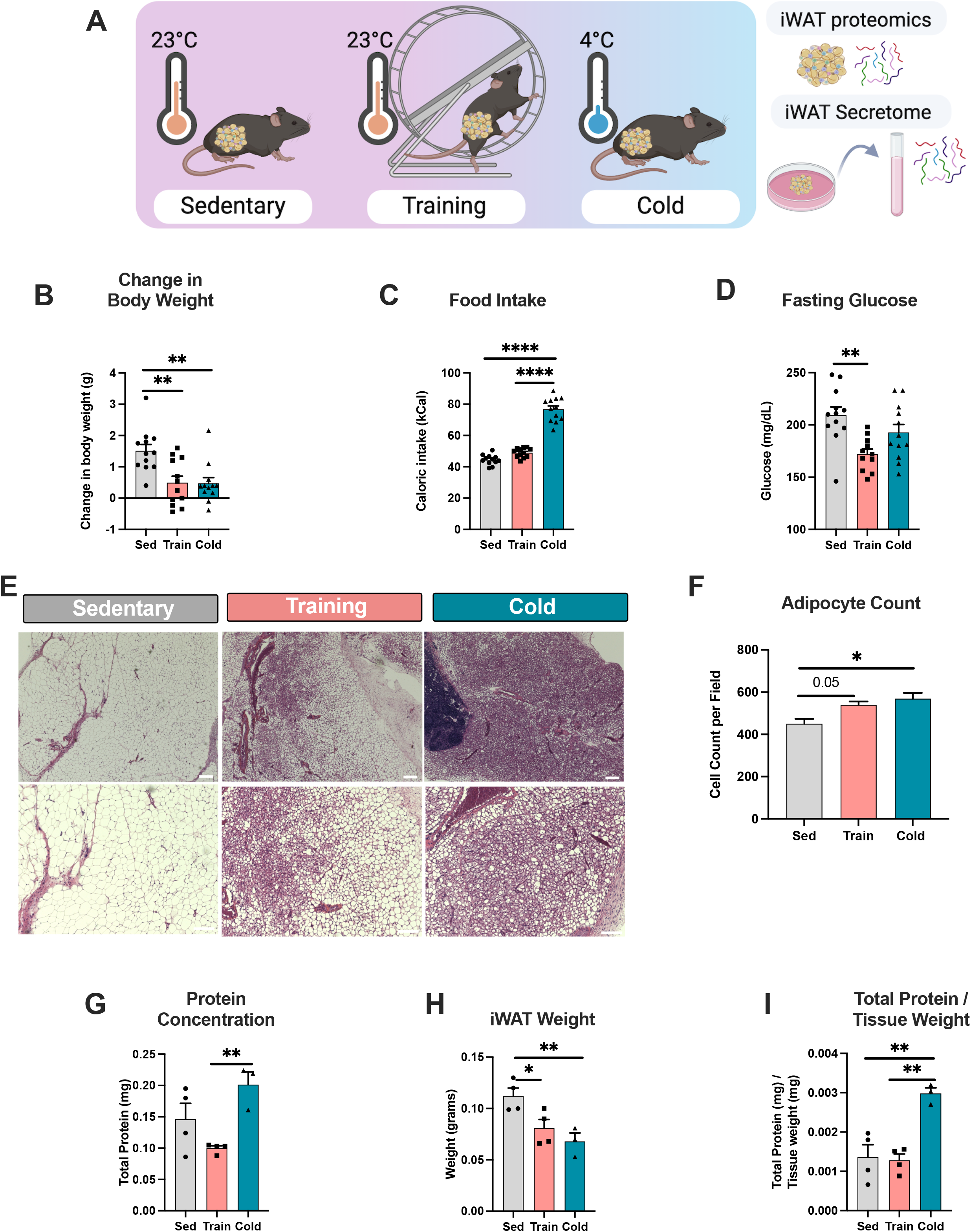
Phenotypic responses of mice and their iWATs to exercise training and cold exposure. (A) Study design to perform quantitative proteomics of whole iWAT and secretome proteins from sedentary, exercise trained and cold-exposed mice. (B-D) Change in body weight (B), food intake (C) and glucose after 6hrs of fasting (D) of sedentary (*gray*), exercise trained (*pink*) and cold-exposed (*green*) mice (*n=12/group*). (E) H&E-stained section of iWAT from sedentary, trained and cold-exposed mice. Scale bar 1000μm (4x, *up*) and 50 μm (20x, *down*). (F) Adipocyte cell count per field measurement of iWAT from sedentary (*gray*), exercise trained (*pink*) and cold-exposed (*green*) mice (*n=5/group*). (G-I) Protein concentration (G), iWAT mass weight (H) and total protein/tissue weight ratio (I) for iWAT from sedentary, exercise trained and cold-exposed mice used for tissue proteomics analysis (*n=3-4/group*). Data are presented as mean±SEM and were compared using One-way ANOVA. *p<0.05,**p<0.01, and ***p<0.001 (E-F) UpSet intersection diagram showing the upregulated (E) and down-regulated (F) proteins compared to sedentary in iWAT secretome from exercise trained (*pink*) and cold-exposed (*green*) mice.

An aliquot of iWAT (∼ 60mg) was evaluated by microscopy and revealed that both exercise training and cold exposure led to significant changes in the appearance of the tissue, including beiging, smaller adipocytes, and an increase in multilocular adipocytes (Figure 2E). However, the effects of cold exposure on these parameters were more pronounced, as was the increase in adipocyte count per field (Figure 2F). For the proteomics, the entire right inguinal fat pad from each mouse was analyzed. Interestingly, the cold-exposed iWAT fat pads had significantly higher protein concentrations than the exercise-trained iWAT fat pads (Figure 2G). Both exercise-trained and cold-exposed iWAT depot weighed less compared to the sedentary iWAT (Figure 2H). When considering the protein concentration per tissue weight, the cold-exposed iWAT demonstrated a higher ratio than both the exercise-trained and sedentary groups (Figure 2I).

### Exercise training and cold exposure effects on iWAT proteome and secretome

To assess the proteome, we collected the entire right inguinal fat pad from 4 sedentary, 4 trained, and 3 cold-exposed mice. Proteomic analysis revealed that exercise training and cold exposure resulted in substantially different effects on iWAT protein expression (Table S1). Principal component analysis (PCA) demonstrated distinct clusters for each treatment group, with the cold exposure group forming a separate and distant cluster from the trained and sedentary groups (Figure 3A). When compared to iWAT from sedentary mice, cold exposure induced robust changes in protein expression, resulting in altered expression of 3459 proteins, whereas exercise training only altered expression of 537 proteins. Cold resulted in the upregulation of 956 proteins (Figure 3B, C), exercise training increased the expression of 300 proteins (Figure 3B), and of these proteins, only 73 proteins were commonly upregulated in both cold-exposed and trained iWAT (Figure 3B). Cold exposure resulted in a striking downregulation in 2096 proteins, while exercise training had a much milder down regulatory effect, with only 182 proteins showing decreased expression (Figure 3C). Interestingly, of these down regulated proteins, 124 were also downregulated in the cold-exposed iWAT, while the remaining 58 were uniquely downregulated by exercise (Figure 3C).

**Figure 3.**
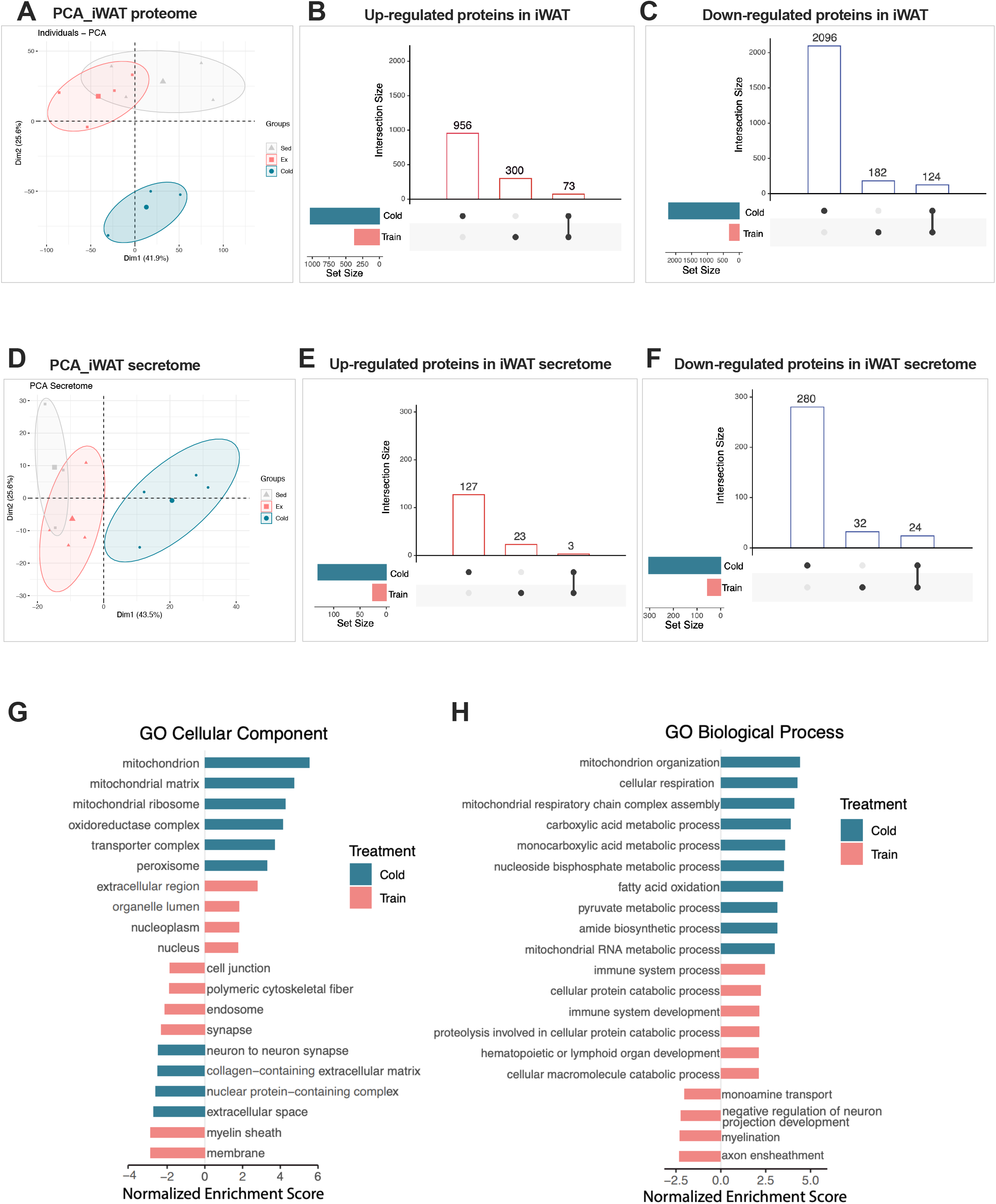
Distinct proteome and secretomic profiles of the exercise and cold-exposed iWATs. (A) PCA plot, including all protein quantified in iWAT from sedentary, exercise trained and cold-exposed mice (*n=4 sedentary*, *4 training*, *3 cold*). (B-C) UpSet intersection diagram showing the upregulated (B) and down-regulated (C) proteins compared to sedentary in iWAT from exercise trained (*pink*) and cold-exposed (*green*) mice. (D) PCA plot, including all protein quantified in iWAT secretome from sedentary, exercise trained and cold-exposed mice (*n=3 sedentary*, *4 training*, *4 cold*). (G) Results of Gene Set Enrichment (GSE) analysis for cellular components of proteins that significantly change in iWAT following exercise training *(pink)* and cold exposure *(green)*. The pathways shown here are considered significant after false discovery correction p < 0.05. (H) Results of Gene Set Enrichment (GSE) analysis for biological process of proteins that significantly change in iWAT following exercise training *(pink)* and cold exposure *(green)*. The pathways shown here are considered significant after false discovery correction p < 0.05.

To investigate the iWAT secretome, iWAT from additional mice (3 sedentary, 4 trained, 4 cold) was used to generate adipose tissue organ culture (ATOC). For the ATOC a small portion of iWAT (20 mg) was incubated with a basic culture medium for a duration of 48 hours. The PCA plot exhibited similar patterns to the tissue proteomic results with all three groups forming distinct clusters. Notably, the cold-exposed group was completely separated from the others (Figure 3D), indicating a clear and unique response to cold exposure. The secretome analysis detected a total of 726 proteins, ∼10.5% of the total number of proteins found in iWAT proteome (Table S2). Cold exposure led to an upregulation of 127 proteins (Figure 3E) while there was a marked downregulation of protein expression, with 280 proteins showing reduced levels (Figure 3F). In contrast, exercise training resulted in a less pronounced change in the iWAT secretome, with only 23 proteins displaying increased expression and 32 showing decreased levels (Figure 3E,F). Our findings highlight the distinct proteomic and secretomic profiles of the exercise and cold-exposed iWATs and the prevalence of downregulated secreted proteins with cold exposure.

### Exercise-training and cold-exposure results in unique adaptations to iWAT cellular components

Using Gene Set Enrichment (GSE) analysis with a focus on Cellular Component (CC) we conducted a comparative analysis of differentially expressed proteins in iWAT to identify the most enriched pathways associated with each treatment. In exercise-trained iWAT there was upregulation of proteins localized in the extracellular region, organelle lumen, nucleoplasm, and nucleus (Figure 3G). The most enriched upregulated pathway was the extracellular region, suggesting that exercise training results in an increased production of proteins secreted into the extracellular matrix potentially enhancing cell-to-cell communication or tissue remodeling (Figure 3G). On the other hand, training resulted in a noticeable downregulation in proteins localized in cell junctions, cytoskeletal fibers, synapses, and the myelin sheath. The reduced focus on maintaining rigid structures (cell junctions, cytoskeleton) could indicate a more dynamic state, perhaps more capable of adapting to changing metabolic needs. In contrast to the trained iWAT, cold-exposed iWAT exhibited a pronounced regulation of numerous aspects of mitochondrial processes (Figure 3G). Cold exposure upregulated proteins in the mitochondrial matrix, mitochondrial ribosome, oxidoreductase complex, transporter complex, and peroxisome, findings suggesting heightened energy expenditure. At the same time, there was a downregulation in neuron-to-neuron synapse, collagen-containing extracellular matrix, nuclear protein-containing complex, and extracellular space with cold. These changes suggest that cold exposure enhances iWAT’s energy-producing capabilities, possibly at the expense of some cellular communication and structure maintenance functions.

### Exercise training and cold exposure have distinct effects on iWAT molecular pathways

The GSE analysis for biological processes showed that exercise training resulted in an upregulation of processes related to cellular protein catabolism, immune system activity, and macromolecule catabolic processes, as well as hematopoietic or lymphoid organ development in iWAT (Figure 3H). These alterations may reflect enhanced protein turnover and immune response, possibly facilitating tissue remodeling and adaptations to exercise-induced stress. Conversely, training resulted in a downregulation in monoamine transport, myelination, axon ensheathment, and negative neurotransmitter level regulation (Figure 3H). This may represent potential changes in neural connectivity, neurotransmitter balance, and neural function following exercise training in iWAT^17,18^. Taken together, these post-exercise alterations indicate that physical activity could stimulate metabolic and immunological adaptations in iWAT, while also modifying its neural interactions. In cold exposure, despite an overall downregulation in protein expression in iWAT as shown in Figure 3C, the majority of significantly enriched pathways were upregulated, with a particular emphasis on mitochondrial-related processes. Cold enhanced the activity of pathways associated with mitochondrial organization, cellular respiration, mitochondrial respiratory chain assembly, carboxylic acid metabolism, fatty acid oxidation, pyruvate metabolic process, and amide biosynthetic process (Figure 3H).

To provide a broad overview of the biological processes and pathways affected by training and cold exposure we performed KEGG pathway analysis. Figure S1A shows the top 10 most enriched pathways with training and cold exposure, again revealing distinct adaptations to iWAT in response to these stimuli. Exercise training upregulated pathways related to calcium signaling, Fc gamma R-mediated phagocytosis, and neurotropin signaling in iWAT and also downregulated pathways such as GnRH signaling, chemokine signaling, GAP function, spliceosome, endocytosis, ERBB pathway, and lysosomes (Figure S1A). These findings suggest that exercise induces significant changes in cellular signaling, immune responses and metabolic pathways within iWAT. In contrast, cold exposure led to upregulation of pathways related to oxidative phosphorylation, peroxisome, aminoacyl-tRNA biosynthesis, valine, leucine, and isoleucine degradation, propanoate metabolism, and fatty acid metabolism (Figure S1A). Upregulation of these pathways suggests that cold exposure prompts iWAT to enhance energy production and thermogenesis through oxidative phosphorylation, lipid and propanoate metabolism, protein synthesis via aminoacyl-tRNA biosynthesis, and an increased reliance on branched-chain amino acids (BCAA) degradation. There was also an upregulation of pathways related to Parkinson’s disease, Alzheimer’s disease, Huntington’s disease suggestive of a potential stress response and neuroprotective mechanisms activated in iWAT to counteract the increased metabolic demands and oxidative stress caused by cold exposure.

Taken together, our GO and KEGG analysis offer insights into iWAT’s distinct physiological responses to exercise and cold exposure. Exercise promotes immune modulation, tissue regeneration, and metabolic regulation in iWAT. In contrast, during cold exposure iWAT adapts with enhanced mitochondrial biogenesis, energy production, and fatty acid metabolism, indicating a more metabolically active state to counteract cold stress. This response to cold, accompanied by increased oxidative stress, prioritizes maintaining body temperature through heightened metabolic processes.

### Exploring iWAT secretome changes post exercise and cold exposure

The exercise-trained iWAT secretome showed a limited number of proteins, precluding detailed pathway analysis. On the other hand, and as illustrated in Figure 3F, cold exposure significantly reduced secreted proteins. These were related to lipid efflux regulation, particularly cholesterol, and the governance of vascular and neuron extension processes (Figure S1B). Cold-upregulated secreted proteins indicated heightened utilization of all macronutrients, such as glucose, lipids, and amino acids as well as increased synthesis of reactive oxygen species, revealing a strong association with the catabolic process (Figure S1C).

To discern important tissue-specific adaptations and systemic responses to exercise and cold exposure we integrated datasets from both the tissue proteome and the secretome and identified upregulated proteins using the Rank Product test. After applying significance and peptide count filters, we identified 17 proteins that were consistently upregulated in both tissue and secretome with exercise training. String analysis revealed two main clusters, one associated with proteasome and antigen presentation, and the other related to amino acid biosynthesis (Figure S2A). The GO analysis of these proteins highlighted their involvement in glutathione and carbohydrate metabolism, positive regulation of protein transport, protein folding, and antigen presentation (Figure S2B). In contrast, cold exposure resulted in the upregulation of 50 proteins in both tissue and secretome. String analysis identified three clusters associated with the TCA cycle, glycolysis/gluconeogenesis, and fatty acid degradation (Figure S2C). The GO analysis of these proteins indicated pathways related to energy oxidation of organic compounds, monocarboxylic acid metabolism, pyruvate metabolism, respiratory electron transport, 2-oxoglutarate metabolism, acetyl-CoA and succinate metabolism, mitochondrial protein localization, and regulation of reactive oxygen species (ROS) (Figure S2D). In exercise training there is a focused response aimed at tissue repair and immune modulation. Conversely, cold exposure induces a more extensive set of upregulated proteins involved in energy metabolism, mitochondrial functions, and the regulation of reactive oxygen species, indicating a broader response to enhance thermogenesis and cope with cold stress.

### Exercise-induced protein signatures in iWAT correlate with improved fasting glucose

Our discovery that mice transplanted with trained, but not cold-exposed iWAT exhibit improved glucose tolerance led us to search for proteins that may be associated with this beneficial effect of training. First, we identified the top 50 proteins for training (Figure S3A) and cold (Figure S3B) that exhibited the most significant condition-specific differential expression. In other words, for exercise, we identified proteins that were most significantly altered compared to the sedentary state and were unaffected by cold, and for cold exposure, we pinpointed proteins most altered compared to sedentary conditions that were not influenced by exercise. We then performed correlation analyses to explore the relationship between these two sets of 50 proteins and fasting glucose levels. In the case of exercise training, remarkably, all 50 proteins exhibited an inverse correlation with fasting glucose levels (Figure 4A, S3C). Conversely, the top 50 differentially expressed proteins in response to cold exposure exhibited a concordant correlation with fasting glucose (Figure 4A, S3D). This proteomic analysis of iWAT in conjunction with the correlation analysis of fasting glucose levels is consistent with our transplantation study findings. The differing correlations between exercise-induced upregulated proteins and enhanced fasting glucose levels emphasize that the remodeling and adaptations occurring in iWAT as a result of exercise are distinct from those triggered by cold exposure.

**Figure 4.**
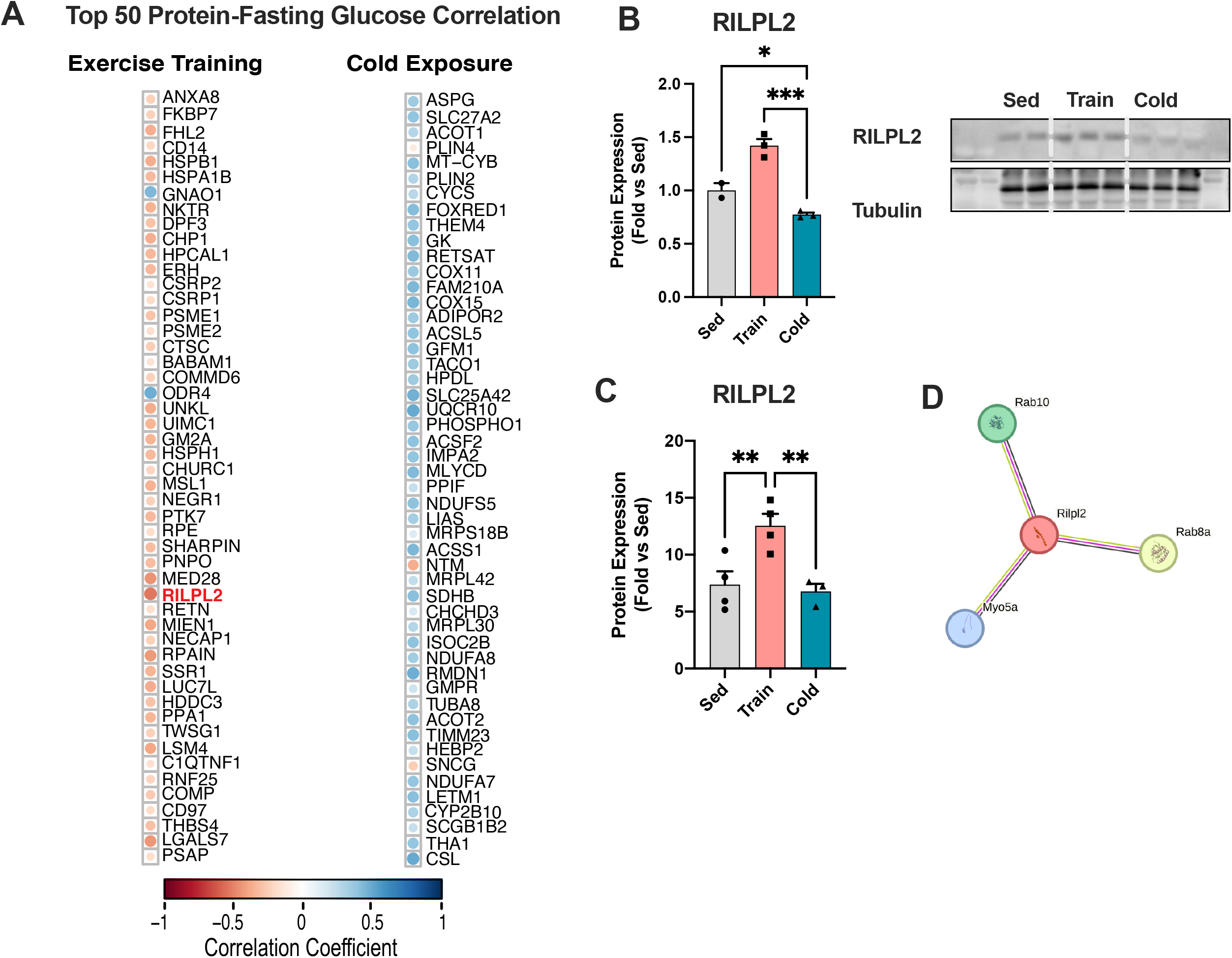
Top 50 differentially expressed proteins in iWAT: Insights into glucose metabolism and Rilpl2’s role in exercise-induced molecular responses. (A) Global correlation matrix of fasting glucose and the top 50 proteins changing in iWAT with exercise training (*left*) and cold exposure (*right*). The color of the circles in the matrix represents the level of correlation for the values that reach statistical significance (P<0.05); blue represents positive correlation and red represents negative correlation. (B) Representative images with relative quantification of Rilpl2 protein detected by western blot in iWAT from sedentary (*gray*), exercise training (*pink*) and cold-exposed (*green*) mice (*n=3-4/group*). (C) Protein abundance level for Rilpl2 detected in the proteome dataset of iWAT from sedentary (*gray*), exercise training (*pink*) and cold-exposed (*green*) mice (*n=3-4/group*) (D) Protein-protein interaction network analysis of Rilpl2 using STRING database with the highest interaction confidence score (0.9) Data are presented as mean±SEM and were compared using One-way ANOVA. *p<0.05, and ***p<0.001

### Rilpl2 is an exercise-induced molecule in iWAT related to glucose metabolism

Considering the significant benefits of exercise-induced iWAT adaptations on glucose metabolism, we specifically delved into the proteins correlated with an improved fasting glucose profile for our study. Notably, Rilpl2 (Rab-interacting lysosomal protein-like 2) from exercise-trained iWAT emerged as a compelling protein, displaying the most pronounced correlation with fasting glucose levels (Figure 4A). Our analysis of the tissue proteome revealed that Rilpl2 levels increased in response to exercise training, but not cold exposure (Figure 4C). This specific effect of exercise on Rilpl2 expression was confirmed by western blot analysis from iWAT of an additional cohort of mice (Figure 4B). To elucidate the potential functional relevance of Rilpl2, we undertook a String analysis solely based on this protein. String analysis helps in predicting protein-protein interactions and can uncover potential associations and co-expression patterns in the context of cellular networks. This analysis unveiled a close association of Rilpl2 with Rab8a, Rab10 and Myosin Va (Myo5a) (Figure 4D), all proteins that are implicated in intracellular transport and vesicle trafficking ^19,20^.

We next investigated the impact of exercise on Rilpl2 expression at the single-cell level using spatial transcriptomics in iWAT. In the sedentary state, Rilpl2 expression levels were low overall in the various cell clusters. However, after 11 days of exercise training, there was a significant increase in Rilpl2 expression in the beige adipocyte cluster (Figure 5A-C). Remarkably, exercise training led to the upregulation of the three proteins closely associated with Rilpl2 (Figure 5D-F). We found a significant exercise-induced increase in expression of Rab8a and Myo5a within the beige adipocyte cluster (Figure 5D-F) while Rab10 expression rose across all cellular clusters post-exercise (Figure 5F). Pathway analysis of this Rilpl2 protein network demonstrates its pivotal role in cellular processes involving protein localization, transport, and secretion, including the cellular response to insulin stimulus, within the context of organelle organization and cell differentiation in the endomembrane system (Figure S4A). Based on these findings, our hypothesis suggests that exercise training influences vesicle trafficking and protein secretion by modifying the expression of key proteins within the Rilpl2-Rab8a-Rab10-Myo5a network with Rilpl2 playing a central role.

**Figure 5.**
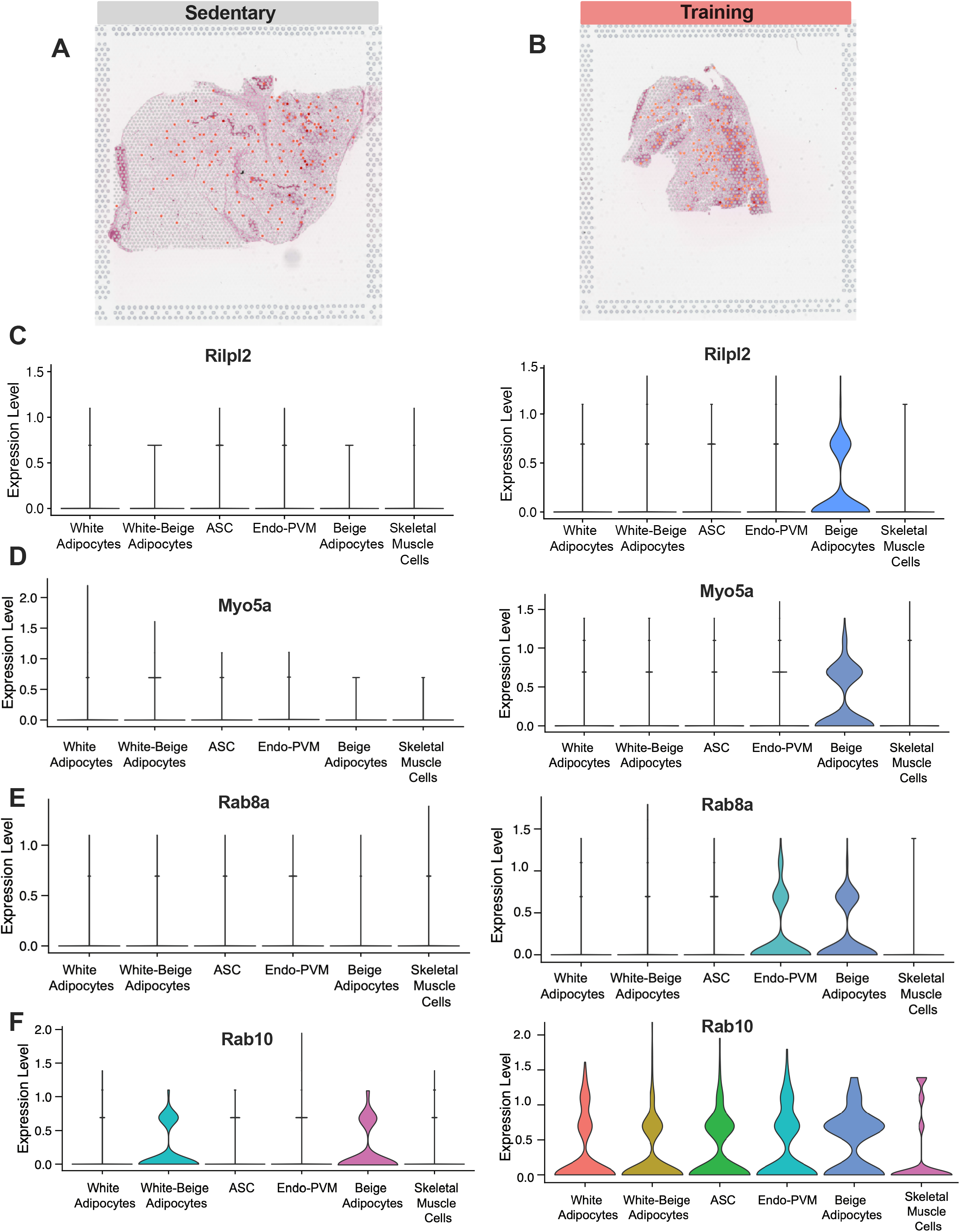
Expression patterns of the Rilpl2 protein network in iWAT in response to exercise training. (A-C) Visium images (A-B) and relative individual violin plots (C) showing the Rilpl2 expression level across the cell clusters detected in iWAT from sedentary (left) and exercise (right). (D-F) Individual violin plots showing the expression levels and distribution of Myo5a (D), Rab8a (E) and Rab10 (F) across cell clusters detected in iWAT from sedentary (left) and exercise (right).

Since spatial transcriptomics primarily identifies cell clusters and not individual cell types, we employed a publicly available single-nuclei dataset from sedentary mouse iWAT to pinpoint the specific cell types associated with Rilpl2 expression^21^. This analysis demonstrated that Rilpl2 is primarily expressed in mature adipocytes and secondary from immune cells such as macrophages (Figure S4B). Upon examination of different adipocyte subpopulations, we discovered that a subpopulation displaying elevated levels of thermogenic genes (mAd5) exhibited the highest expression of Rilpl2 (Figure S4C,D). These findings align with our spatial transcriptomic results, where exercise-induced elevation of Rilpl2 expression was observed in the beige cluster. To gain deeper insights into the regulation of Rilpl2 expression at the single-cell level in response to exercise training, we leveraged our recently published single-cell dataset, encompassing iWAT sc-RNAseq data from both sedentary and trained mice. This approach, while not primarily focused on mature adipocytes, offers a distinctive perspective by shedding light on stem cells and specific immune cell populations ^6^. Using this dataset we identify Rilpl2 expression in immune cells including macrophages, monocytes, dendritic cells, plasma cells, and B cells (Figure S4E,F). When determining expression levels of Rilpl2 with all cell types studied combined, we found that exercise training upregulates the number of cells expressing Rilpl2 (Figure S4G). Taken together, our findings demonstrate that Rilpl2 expression is specifically upregulated by exercise training in adipocytes with heightened thermogenic capacity and immune cells, suggesting it may play a role in exercise-induced intracellular transport and vesicle trafficking orchestrating appropriate protein distribution and secretion.

### Exercise training increased the production and release of small extracellular vesicles from mature adipocytes (Ad-EVs)

Despite the robust systemic effects of exercise-induced adaptations in iWAT, our analysis revealed a relatively small number of secreted proteins within the iWAT secretome. This discovery raises intriguing questions regarding the possibility that iWAT might exert systemic effects through the secretion of essential proteins encapsulated within vesicles. Additionally, our pathway analysis and data suggest that exercise training induces notable changes in intracellular transport and vesicle trafficking, crucial processes in regulating protein distribution and secretion. In light of these findings, we have formulated a novel hypothesis proposing that exercise training triggers the trafficking and secretion of small extracellular vesicles (Ad-EVs) originating from adipocytes in iWAT. We used size exclusion chromatography (SEC) to investigate if mature adipocytes, freshly isolated from sedentary and trained mice, release Ad-EVs into the conditioned culture media (Figure 6A). Following isolation of the Ad-EVs, we performed quality control using transmission electron microscopy (TEM) and nanoparticle tracking analysis (NTA). We found that the Ad-EVs, which were positive for the exosomal marker CD63, measured between 50 and 200 nm in diameter (Figure 6B). Mature adipocytes from the exercise-trained mice produced and released significantly more Ad-EVs than those from the sedentary group, despite there being no significant differences in the size of the Ad-EVs between the two groups (Figure 6C and D). These data provide evidence that exercise training leads to a substantial increase in the production and release of Ad-EVs. We hypothesize that these exercise-triggered Ad-EVs may have a substantial impact on facilitating the positive outcomes of physical activity by fostering communication between different tissues.

**Figure 6.**
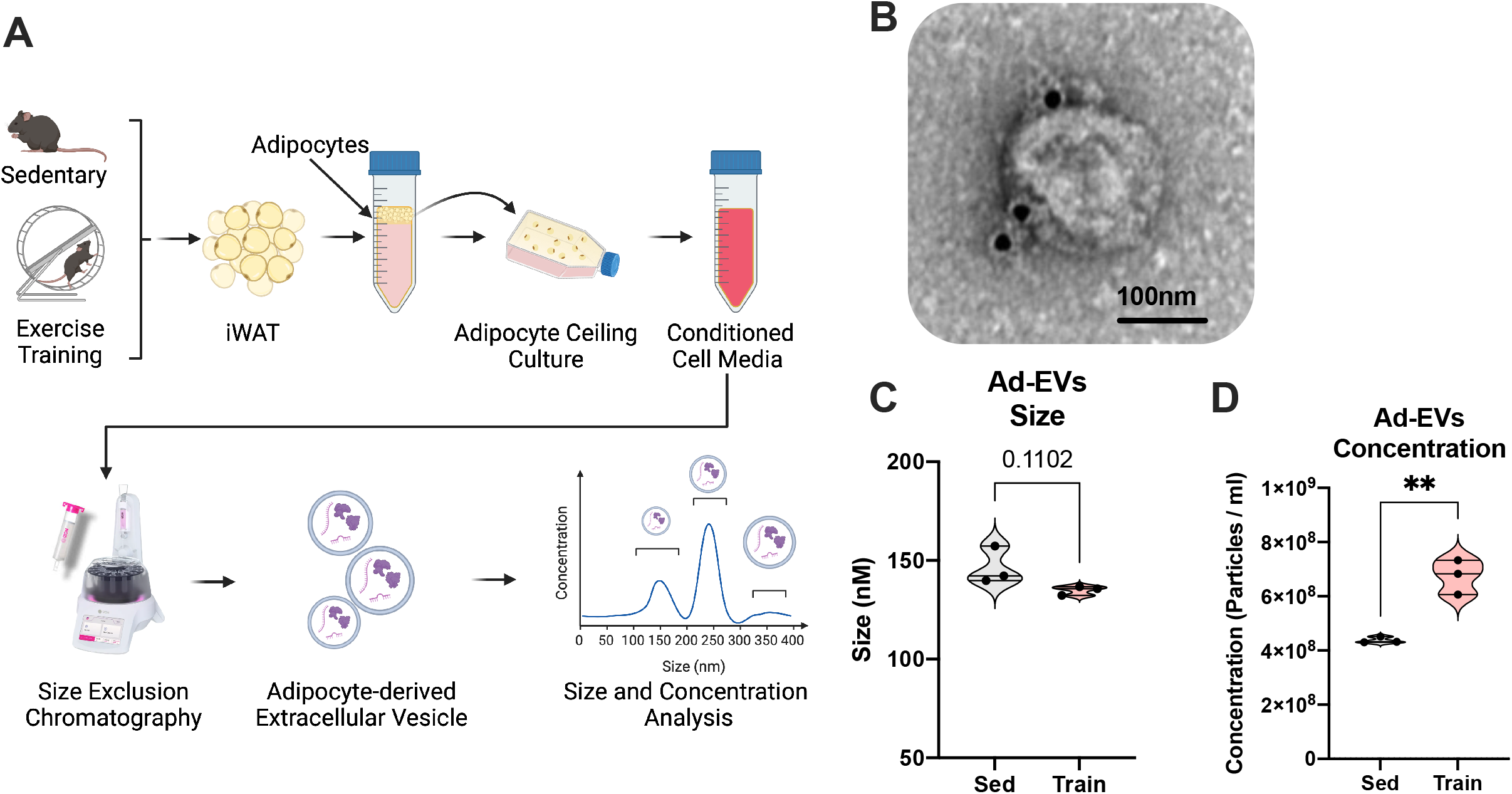
Adipocyte-derived extracellular vesicles secretion in response to exercise training. (A) Study design to isolate Adipocyte-derived extracellular vesicles (Ad-EVs) from fresh isolated mature adipocytes from iWAT of sedentary and trained mice, using the size exclusion chromatography (SEC). (B) Representative electron micrograph showing CD63 gold immunostaining (1:20, 10nm) of Ad-EVs isolated from mature adipocytes using SEC. Scale bar: 100nM. (C-D) Nanoparticle Tracking Analysis (NTA) of Ad-EVs isolated from sedentary and trained iWAT (n=3 biological replicates, from two Ad-EVs isolation) representing the size (C) in nm and the number of particles per mL (C) of adipocyte suspension. Data are presented as mean±SEM and were compared using unpaired two-tailed Student’s t-test. **p<0.01.

## DISCUSSION

This study unveils dramatic distinctions in the effects of exercise training and cold exposure on iWAT, shedding light on their individual proteomic profiles. While both trained and cold-exposed iWAT exhibit some similar phenotypic changes, including reduced adipocyte size and increased beiging, they significantly diverge in their systemic impact on glucose metabolism, as evidenced by the transplantation experiments. Proteomic analysis reveals the role of exercise training in inducing tissue remodeling, modulating immune responses, and upregulating protein expression in iWAT. In contrast, cold exposure primarily downregulates protein synthesis while enhancing processes related to mitochondrial protein synthesis and thermogenesis. Exercise training results in an increase in the expression of cytosolic proteins involved in cellular processes related to protein secretion, exocytosis, and vesicle trafficking, suggesting exercise’s role in enhancing the endocrine function of iWAT. Importantly, the proteins exclusively influenced by exercise training are associated with a favorable glycemic profile, consistent with our transplantation studies, thereby distinguishing exercise-induced adaptations from those induced by cold exposure.

A notable finding of our study is that cold-exposed iWAT, despite showing profound beiging, did not yield any systemic benefits in terms of glucose metabolism when transplanted into sedentary recipient mice. While we did identify a prominent number of upregulated proteins in cold-exposed iWAT, these were closely related to processes such as oxidative phosphorylation, peroxisome pathways, and branched-chain amino acid degradation, processes well-established in the realm of thermogenesis^22^. What truly stood out was the striking downregulation of nearly 2100 proteins. Other studies have reported a noticeable decrease in the expression of adipokines and cytokines in cold-exposed iWAT ^9,23,24^. Here, we found that these down regulated proteins were associated with pathways linked to cholesterol efflux and extracellular components, suggesting a potential disruption of the endocrine function of the cold-exposed iWAT ^25,26^. Our data confirm the pivotal role of cold exposure in promoting thermogenesis in iWAT, effectively transforming this adipose depot into a potent energy-generating powerhouse ^27,28^. However, our proteomic analysis along with the iWAT transplantations findings underscore the limited impact of this transformation on enhancing glucose metabolism. These results imply that improvements in glucose metabolism following cold exposure may primarily be attributable to adaptations in BAT and not iWAT, since BAT has been shown to have a central role in improving glucose metabolism in response to cold temperatures ^29–32^.

Exercise training resulted in significant phenotypic changes in iWAT, akin to those induced by cold exposure, such as the presence of smaller adipocytes and the emergence of beiging. However, a key distinction emerged when we transplanted trained iWAT into sedentary recipients: unlike cold-exposed iWAT, the trained iWAT significantly improved glucose metabolism. This outcome strongly implies systemic effects of the trained iWAT and an enhanced endocrine function. Although the number of proteins regulated by exercise training was lower than in cold exposure, our proteomic analysis uncovered distinct patterns. Upregulated proteins in trained iWAT were associated with tissue remodeling, immunomodulation, and the extracellular space. They were also found to be involved in the structuring of organelle interiors and vesicle transport. These findings suggest that exercise enhances cell-to-cell communication by optimizing intracellular movement, protein maturation and release ^33^. The exercise-induced downregulation of proteins linked to cell junctions and cytoskeletal fiber implies a decrease in cell adhesion and potentially promotion of increased cell mobility within the iWAT, and this may further enhance the exercise-induced cell-to-cell communication. It is also possible that these cytoskeletal changes are associated with the known reduction in adipocyte size, leading to decreased requirements for cytoskeleton protein synthesis^34^.

An intriguing contrast arose when comparing the tissue proteome with the secretome analysis of trained iWAT. While the tissue proteome hinted at an enhanced metabolic and endocrine function in trained iWAT, the secretome analysis revealed a more limited number of upregulated proteins in the trained group, whereas cold exposure resulted in more substantial protein alterations. Building upon our tissue proteome analysis, which suggested increased exercise-induced vesicle trafficking and protein exocytosis, we proposed that exercise training might induce the secretion of proteins encapsulated within secreted vesicles which we failed to detect with the method employed for our secretome analysis. In fact, adipose tissue has recently emerged as a major source of circulating small extracellular vesicles (EVs) or exosomes with a potentially significant role in promoting interorgan communication and affecting metabolic health^35^. To explore this hypothesis, we isolated adipocyte-derived EVs (Ad-EV) released from sedentary and trained mice and found a significant increase in EV release from the exercise trained adipocytes. This discovery reinforces the concept that exercise training facilitates the release of iWAT-derived molecules, fostering inter-tissue communication. Furthermore, it establishes a foundation for future research into the role of Ad-EVs within the context of exercise.

The substantial impact of the exercise-induced iWAT adaptations on overall glucose metabolism motivated us to investigate particular proteins that could be pivotal in driving these exercise-induced metabolic alterations. Rilpl2 stood out due to its robust correlation with fasting glucose levels, a measurement intricately linked to metabolic health. The potential role of Rilpl2 in regulating glucose homeostasis has been supported by the elevated fasting glucose levels in a Rilpl2 knockout mouse model ^36^. Rilpl2 is recognized for its involvement in cellular glycolytic processes in cancer cells ^37^ while studies in neuronal cells and adipocytes have demonstrated its interaction with Myosin Va ^38^, an integral part of the Rac1-Akt2 pathway critical for insulin-facilitated GLUT4 translocation and glucose intake ^39,40^. Here, we showed that Myosin Va expression is concomitantly increased with exercise training in the same cell cluster as Rilpl2, supporting a possible interaction of these molecules in the context of exercise and their potential role for the Rilpl2-Myosin Va axis in regulating glucose transport. Additionally, Rilpl2 is associated with various Rab proteins which are known to be significant regulators of vesicular transport^41^. Rab8a and Rab10 are known to be the main interacting proteins with Rilpl2 and both were found to have similar increased expression patterns following exercise training. These findings may indicate the Rilpl2’s indirect influence on vesicular trafficking. Such multifaceted potential positioning of Rilpl2 offers intriguing directions for future research into this protein’s role in glucose homeostasis. It is plausible that Rilpl2 might indirectly contribute to Ad-EVs generation and/or trafficking through its interactions with the Rab GTPases and thus, further studies will be needed to determine the role of Rilpl2 in Ad-EVs biogenesis, trafficking, and secretion.

In conclusion, despite exercise training and cold exposure both improving systemic metabolism, our study reveals distinct effects of these physiological stimuli on iWAT, including only trained iWAT having direct effects on in vivo glucose metabolism. Cold-exposed iWAT adapts with enhanced mitochondrial biogenesis, energy production, and fatty acid metabolism in an attempt to counteract cold stress. Conversely, exercise induces iWAT remodeling at both the cellular and extracellular realms, priming the trained iWAT to readily adapt to fluctuating metabolic needs. Our findings hint at a heightened endocrine role of the trained tissue, potentially mediated through increased secretion of adipocyte derived extracellular vesicles.

### LIMITATIONS

All mouse experiments were performed in male mice. We used only male mice for these experiments due to the practical difficulty of handling a large number of mice simultaneously, especially during transplantation experiments. The effects of exercise and cold exposure on female iWAT will be investigated in future studies. The lower number of proteins detected in the exercise-induced iWAT secretome can be attributed to factors such as the potential presence of exercise-induced proteins with lower molecular weights, falling below the detection limits of our mass spectrometry method. Lastly, we identified exercise-induced proteins that were negatively correlated with fasting glucose levels. However, we did not investigate the specific mechanisms through which these proteins impact glucose metabolism. This aspect falls beyond the scope of our current study and will be addressed in future experiments.

## STAR⋆ METHODS

### KEY RESOURCES TABLE

### CONTACT FOR REAGENT AND RESOURCE SHARING

### EXPERIMENTAL MODEL AND SUBJECT DETAILS

Mouse Models

### METHOD DETAILS

Subcutaneous inguinal white adipose tissue transplantation to recipient sedentary mice

Glucose tolerance test in recipient mice

Orbitrap-Based LC-MS/MS Analysis

Quantitative proteomics of secretome proteins from adipose tissue organ culture.

Western Blot

Spatial transcriptomics

Adipocyte-derived extracellular vesicles (Ad-EVs) isolation

Computational analysis

Statistical Analysis

### CONTACT FOR REAGENT AND RESOURCE SHARING

Requests for reagents and resources should be directed to the Lead Contact, Laurie J. Goodyear.

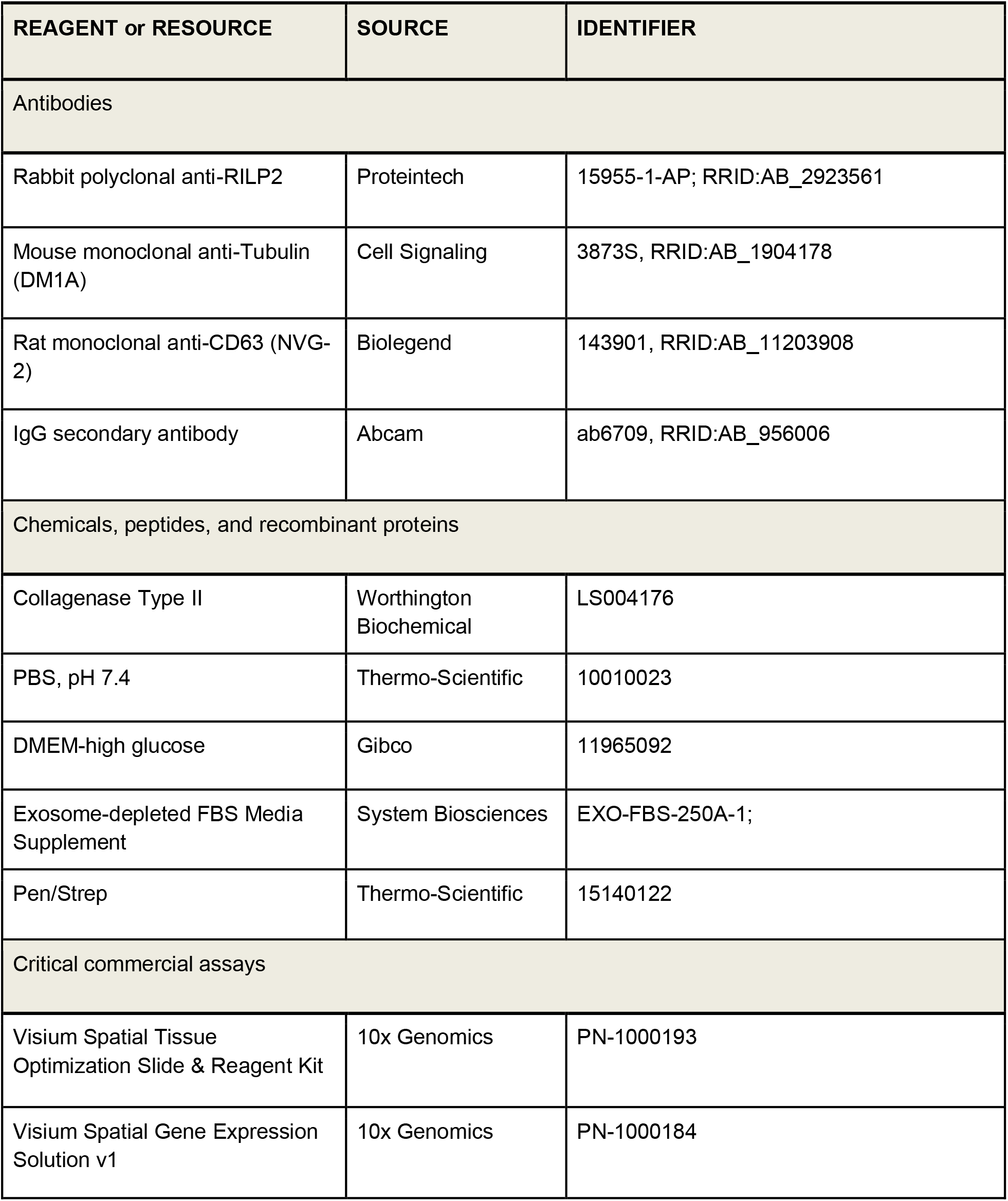
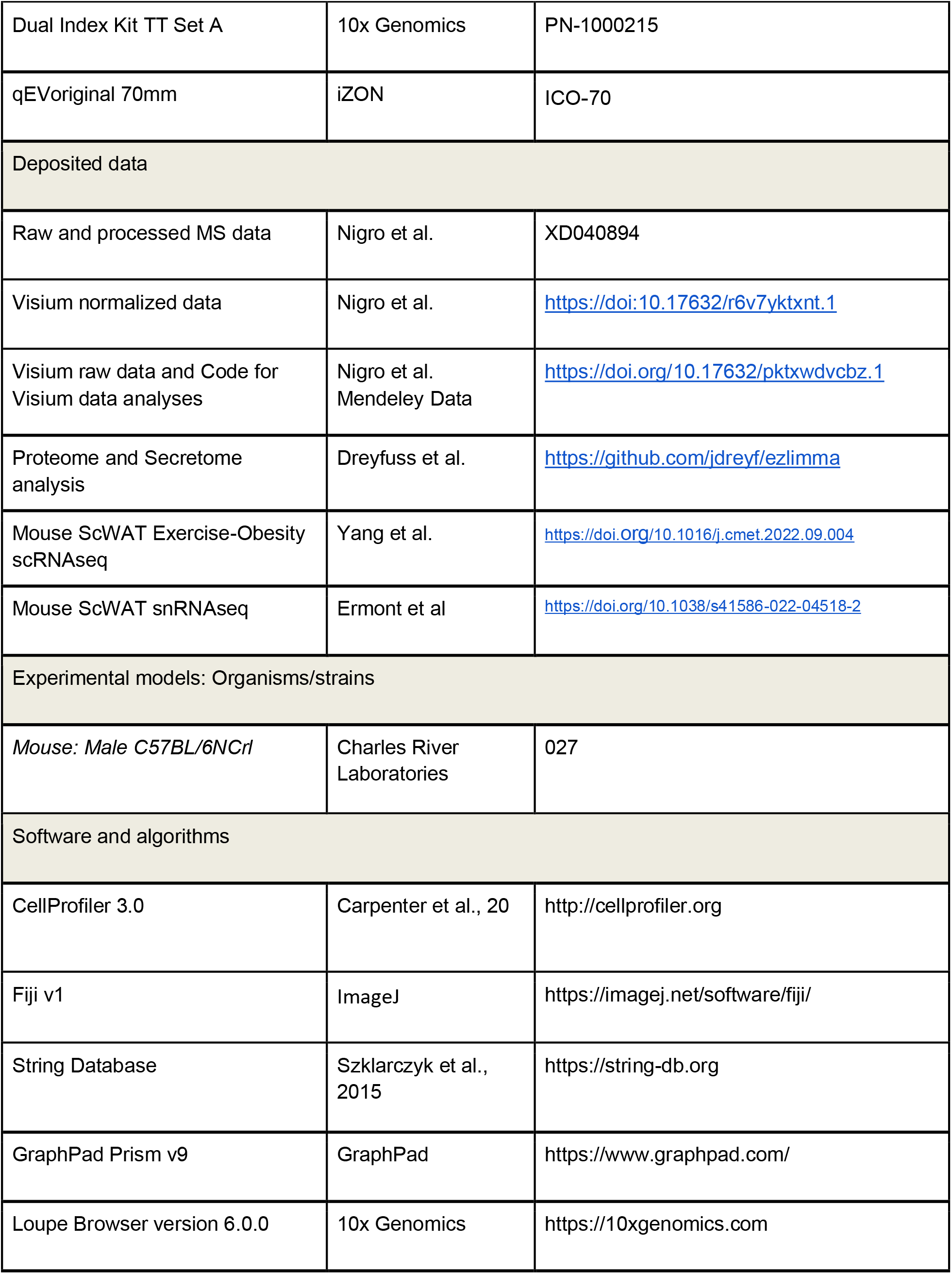
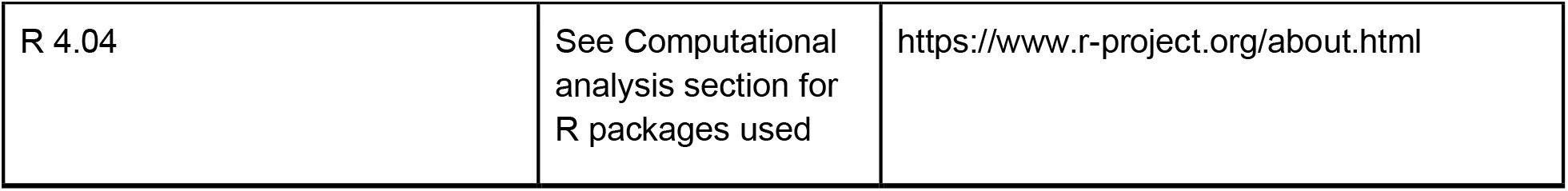

## EXPERIMENTAL MODEL AND METHOD DETAILS

### Mouse 1Models

All experiments were conducted following NIH guidelines and protocols approved by the IACUC at Joslin Diabetes Center (JDC). All experiments were performed using 8-week-old male C57BL/6 mice purchased by Charles River Laboratories, MA. All mice from all groups were housed on a 12/12 h light/dark cycle. Standard mouse chow diet (9F 5020 Lab Diet, PharmaServ, Inc.) and water were provided ad libitum. The group of sedentary and exercise training mice were housed at room temperature at 23°C. For the exercise training experiments, mice had access to a running wheel at all times and performed voluntary running (24 cm diameter; Tecniplast) for 11 days. The total number of wheel cage revolutions was monitored on a daily basis and the accumulated running distance was calculated at the end of the 11 day period (>5 km/mouse). The sedentary mice were housed in similar individual cages without having access to a training wheel. For chronic cold exposure, C57BL/6n mice were individually housed at 5°C for 11 days using temperature controlled diurnal incubators (Caron Products & Services Inc.) on a 12 hour light/dark cycle with ad libitum access to food and water. After 11 days, mice were removed from their cages. For the exercise training group, the wheels were locked 24hrs prior to this. The mice were anesthetized, heart puncture for blood collection was performed and iWAT was harvested immediately. The iWAT was either snap frozen for protein extraction and proteomic analysis or used fresh for adipose tissue organ culture and subsequent adipose tissue secretome analysis or used for transplantation to recipient age-matched mice.

### Subcutaneous inguinal white adipose tissue transplantation experiment

We conducted fat transplantation using WAT obtained from the inguinal area of mice exposed to exercise, cold, or of sedentary conditions. The donor mice were humanely euthanized using heart puncture and cervical dislocation. We carefully removed and kept their fat pads in 1x PBS solution at 37°C until the transplantation procedure. To prepare the recipient mice, we administered intraperitoneal injections of pentobarbital at a dose of 85-100 mg/kg to induce anesthesia. Each recipient mouse received a transplantation of 0.85 grams of subcutaneous WAT into their visceral cavity. We placed this tissue deep within their existing perigonadal adipose tissue, adjacent to the mesenteric adipose tissue just below the liver. To ensure an equal amount of transplanted adipose tissue, approximately four trained mice and two sedentary mice were used for each transplantation. For the control group of recipient mice, we performed a sham procedure, following the same surgical steps without actually transplanting any adipose tissue. Throughout the study, all recipient mice were kept in sedentary conditions in stationary cages. We closely monitored their recovery after surgery and observed no discernible differences in their recovery rates across the groups, as they all had similar body weights and were carefully observed for several days post-transplantation.

### Orbitrap-Based LC-MS/MS subcutaneous adipose tissue proteomic analysis

The peptides were subjected to separation using a gradient ranging from 3% to 36% Buffer B (composed of 90% acetonitrile in 0.1% formic acid) over a period of 120 minutes. The MS analysis was performed using an EASY-nLC™ 1200 System interfaced with an Orbitrap FusionTM TribridTM Mass Spectrometer (Thermo Scientific, United States). For the 11-plex data search using Sequest, the following parameters were employed: Peptide Mass Tolerance = 20 ppm, Fragment Ion Tolerance = 1, Max Internal Cleavage Site = 2, and Max differential/Sites = 4. Regarding the Reported Quant Parameter, the settings were as follows: tolerance = 0.003, ms3 = 1, peak_picking = max, num_isotopes = 2, ms2_isolation_width = 0.7, and ms3_isolation_width = 1.2. The MS2 spectra were searched using the SEQUEST algorithm against a composite database derived from the Mouse proteome available in Uniprot, which included the reversed complement of the sequences and known contaminants. To ensure data quality, peptide spectral matches were filtered to achieve a false discovery rate (FDR) of 1% using a target-decoy strategy in combination with linear discriminant analysis. Additionally, the proteins were filtered to maintain an FDR of less than 1%. Protein quantification was performed exclusively on peptides with a summed signal-to-noise (SN) threshold greater than 200 and an MS2 isolation specificity of 0.5.The mass spectrometry proteomics data were previously deposited to the ProteomeXchange Consortium via the PRIDE partner repository with the dataset identifier PXD040894 ^4^.

### Quantitative proteomics of secretome proteins from adipose tissue organ culture

As previously described^4^, for the analysis of the secretome, we employed Quantitative Immunoprecipitation (Quant-IP) analysis. To begin, 20 mg of inguinal white adipose tissue (iWAT) was subjected to three washes with 1X phosphate-buffered saline (PBS). Subsequently, the tissue was incubated in M199 media supplemented with 1 µM insulin (Sigma, 11376497001), 250 nM dexamethasone (Sigma-Aldrich, D1756), and 1% penicillin/streptomycin (Thermo Scientific, 15140122). Following a 24-hour incubation period, the media was replaced with fresh M199 media containing 1% penicillin/streptomycin. After another 24 hours, the media was collected and immediately stored at -80°C. Debris was eliminated from the media through centrifugation at 4000 × g, and the resulting supernatant containing proteins was combined with a lysis buffer consisting of 50 mM EPPS (pH 8), 1 M urea, 1% SDS, and protease inhibitors (1 mini tablet of Roche EDTA-free protease inhibitor per 10 ml). The tryptic peptides were subjected to reduction and alkylation using a combination of 5 mM TCEP, 15 mM iodoacetamide, and 10 mM DTT at 56°C for 30 minutes. Protein precipitation was achieved by employing TCA, and the resulting precipitate was dissolved in 50 mM EPPS. Trypsin digestion was carried out at a protease-to-protein volume ratio of 1:100. Each sample was labeled with TMT tags for multiplex proteomics analysis.

### Spatial transcriptomics

We utilized the 10x Genomics Visium Spatial Gene Expression platform (10x Genomics, Pleasanton, CA, USA) for our spatial transcriptomics analysis. As previously described, the experiment involved the utilization of paraffin-embedded inguinal white adipose tissue (iWAT) obtained from sedentary and trained mice^4^. The Visium Spatial Gene Expression procedure was conducted following the manufacturer’s instructions (10x Genomics, Pleasanton, CA, USA) with the following parameters: adipose tissue sections of 20 μm thickness were used, and hematoxylin and eosin (H&E) staining was performed by immersing the slides in isopropanol for 1 minute, followed by a 3-minute incubation in hematoxylin and a 30-second incubation in eosin (diluted at 1:20). Permeabilization of the tissue was carried out for 15 minutes. All Visium cDNA libraries were indexed, pooled, and subsequently sequenced simultaneously using the Illumina NovaSeq6000 platform, with support from the BioMicro Center Core at MIT. Following the Visium protocol, the sequencing entailed 28 nt for R1, 120 nt for R2, and 10 nt for each of the indexes.

### Adipocyte-derived Extracellular Vesicles (Ad-EVs) isolation from murine adipocyte iWAT

Adipocyte-derived extracellular vesicles (Ad-EVs) were isolated from the adipocyte fraction of the subcutaneous inguinal adipose tissue (iWAT) obtained from male C57BL/6n mice that underwent either 11 days of voluntary wheel running or remained in a sedentary state. We first processed the iWAT to obtain the mature adipocyte fraction, following a previously described protocol ^42^. Subsequently, the adipocytes were cultured for one week in a flask containing DMEM-high glucose supplemented with 1% Pen/strep and 20% of Exosome-depleted FBS Media Supplement (EXO-FBS-250A-1). The conditioned cell media were subsequently collected, concentrated, and utilized to isolate Ad-EVs through size exclusion chromatography (SEC) using a qEV original 70mm column (iZON, ICO-70), as previously reported^43^. An aliquot of each Ad-EVs sample was used for determining particle concentration through nanoparticle tracking analysis (NTA) and for Transmission Electron Microscopy (TEM). Additionally, CD63 immunogold staining of Ad-EVs preparations was performed at the Electron Microscopy Facility at Harvard Medical School (Boston, MA), following a previously described procedure ^43^.

### Computational analysis

Proteome and secretome scaled relative intensities were processed as follows. For the proteomics data we imputed missing values (zeros) using a random forest algorithm in the R package missForest. We unbiasedly estimated the empirical sample quality weights in the R package limma. We observed a trend between a protein’s average abundance and its variance and accounted for it in variance estimation using limma-trend. For the proteome, we performed Surrogate Variables Analysis (SVA) to remove unknown batch effects (i.e. the unwanted variation that is independent of the group assignment) using the R package sva. We tested the differential abundance of each protein using limma ^44^. Volcano plots were generated using the R package ggplot2. We performed pathway analysis on differentially abundant proteins (P-value < 0.05) using the R package gseGO.

### Statistical Analysis

Data are expressed as mean ± SEM. Sample sizes are indicated in the figure legends. All statistical analyses were performed using GraphPad Prism v9. Statistical significance was analyzed by One-way ANOVA and two-way ANOVA followed by Tukey’s multiple comparisons test or unpaired two-tailed Student’s t-test.

## ACKNOWLEDGMENTS

This work was supported by NIH grant R01DK099511 and R01DK101043 (to L.J.G.), T32DK00726042, F32DK12643201 and Joslin DRC (P30 DK36836) Pilot and Feasibility Program award (to M.V.), K23DK114550 (to R.J.W.M), and the Joslin Diabetes Center DRC (P30 DK36836). We thank Drs Hui Pan and Jonathan M. Dreyfuss from Joslin Diabetes Center for statistical analysis support, Dr. Marsel Lino from Joslin Diabetes Center for his assistance for the extracellular vesicle isolation and Dr. Maria Ericsson from Harvard Medical School for the electron microscopy analysis.

## AUTHOR CONTRIBUTIONS

M.V. and PN designed research, carried out experiments, analyzed data and wrote the manuscript. K.S. carried out experiments. T.C performed bioinformatics analysis. M.F.H. supervised all experiments. R.J.W.M. provided scientific advice and edited the manuscript. L.J.G. directed the research project, designed experiments and wrote the manuscript. All authors have participated in the manuscript review and approved the final manuscript.

## SUPPLEMENTARY FIGURE LEGENDS

**Figure S1 The KEGG pathways analysis supported the findings of the pathway analysis conducted on iWAT samples from exercise-trained and cold-exposed mice. Additionally, the GSE analysis revealed significant changes in the iWAT secretome for both upregulated and downregulated proteins following cold exposure, related to Figure 2**

(A) Summary of the top 10 KEGG pathways changing with exercise training and cold exposure in iWAT compared to sedentary mice.

(B-C) Dot plots for the top 10 Biological pathways down-regulated (A) and upregulated (B) in iWAT secretome following cold exposure. The x-axis shows the ratio of differentially expressed genes in each term relative to the total number of genes in that term. Dot size shows the number of differentially expressed genes in that term and color is the gene set enrichment analysis test statistic with Benjamini-Hochberg adjustment for multiple testing

**Figure S2 The STRING Network Analysis and the corresponding GO Biological Process for integrating proteome and secretome datasets, related to Figure 3**

(A-B) The STRING network (A), employing the default k-means clustering method, displays the three main protein clusters and relative top GO Biological Process Pathways (B) for the integrated proteome and secretome datasets of exercise-trained mice.

(C-D) The STRING network (C) and relative top GO Biological Process Pathways for integrating proteome and secretome datasets of cold exposed (D, *green*) mice.

**Figure S3 List of highly differentially expressed proteins in iWAT following exercise training and cold exposure in mice, related to Figure 4**)

(A-B) Heat maps for the top 50 differentially expressed proteins in iWAT from exercise training (A) and cold exposed mice (B). Rows represent the top 50 proteins, and columns represent the iWATs with their replicates. The colors follow the z-scores (blue low, white intermediate, red high).

(C-D) Extended version for the global correlation matrix of fasting glucose and the top 50 differentially expressed proteins in iWAT with exercise training (C) and cold exposure (D). The color of the circles in the matrix represent the level of correlation for the values that reach statistical significance (P<0.05); blue represents positive correlation and red represents negative correlation.

**Figure S4 Exploring Rilpl2: GO Biological Processes of the Rilpl2 protein network, Cell-Type Expression, and Adipocyte Subclusters in iWAT, related to Figure 4**

(A) Overview of GO Biological Processes Linked to the Rilpl2 Protein Interaction Network in STRING Database.

(B-C) Individual violin plots showing the expression levels and distribution of Rilpl2 across different cell types (B) and specific mature adipocyte subclusters annotated (C) in ScWAT from the GSE176171 dataset.

(D) Annotation analysis for beige/thermogenic mature adipocytes using the Single Cell Portal (https://singlecell.broadinstitute.org/single_cell) and relative Rilpl2 gene expression.

(E-F) Rilpl2 gene expression across the different cell types annotated in scRNA seq from iWAT (GSE183288) represented on a UMAP plot (E) and individual violin plots (F).

(G) Rilpl2 gene expression across the entire iWAT single-cell transcriptomic dataset was assessed using the GSE183288 dataset, encompassing both sedentary and trained mice.

**Table S1: Proteins that were found to be differentially expressed in mouse iWAT following exercise and cold exposure.**

**Table S2: Proteins that that were found to be differentially expressed in the secretome of mouse iWAT following exercise and cold exposure**

