## Supplemental Figure 1 for "Exercise Training and Cold Exposure Trigger Distinct Molecular Adaptations to Inguinal White Adipose Tissue"

**A**

| | KEGG Pathways | Compared to Sedentary | p-value | FDR |
|---|---|---|---|---|
| Exercise Training | GNRH signaling pathway | Down | 3.7E-09 | 1.2E-06 |
| | Chemokine signaling pathway | Down | 4.7E-09 | 1.2E-06 |
| | Gap junction | Down | 6.6E-09 | 1.2E-06 |
| | Spliceosome | Down | 9.5E-08 | 1.2E-05 |
| | Endocytosis | Down | 2.2E-07 | 1.9E-05 |
| | Calcium signaling pathway | Up | 8.2E-07 | 5.7E-05 |
| | Fc gamma R mediated phagocytosis | Up | 1.2E-06 | 7.9E-05 |
| | Neurotrophin signaling pathway | Up | 1.3E-06 | 7.9E-05 |
| | ERBB signaling pathway | Down | 2.0E-06 | 1.1E-04 |
| | Lysosome | Down | 2.9E-06 | 1.4E-04 |
| Cold Exposure | Oxidative phosphorylation | Up | 7.2E-34 | 3.0E-31 |
| | Parkinsons' disease | Up | 3.9E-31 | 9.8E-29 |
| | Huntingtons disease | Up | 5.4E-18 | 9.8E-16 |
| | Alzheimers disease | Up | 6.2E-18 | 9.8E-16 |
| | Valine,leucine and isoleucine degradation | Up | 1.4E-17 | 1.9E-15 |
| | Citrate cycle TCA cycle | Up | 1.2E-14 | 1.3E-12 |
| | Peroxisome | Up | 1.3E-14 | 1.3E-12 |
| | Amino acyl tRNA biosynthesis | Up | 1.4E-14 | 1.3E-12 |
| | Propanoate metabolism | Up | 3.7E-13 | 2.7E-11 |
| | Fatty acid metabolism | Up | 2.9E-12 | 1.8E-10 |

**B**

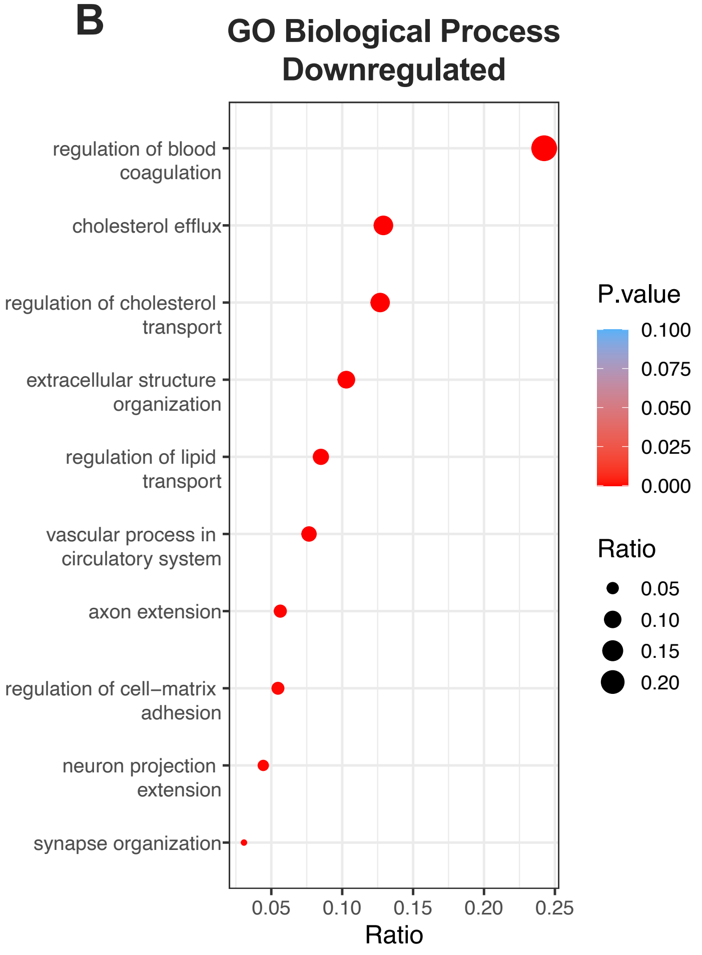


**C**

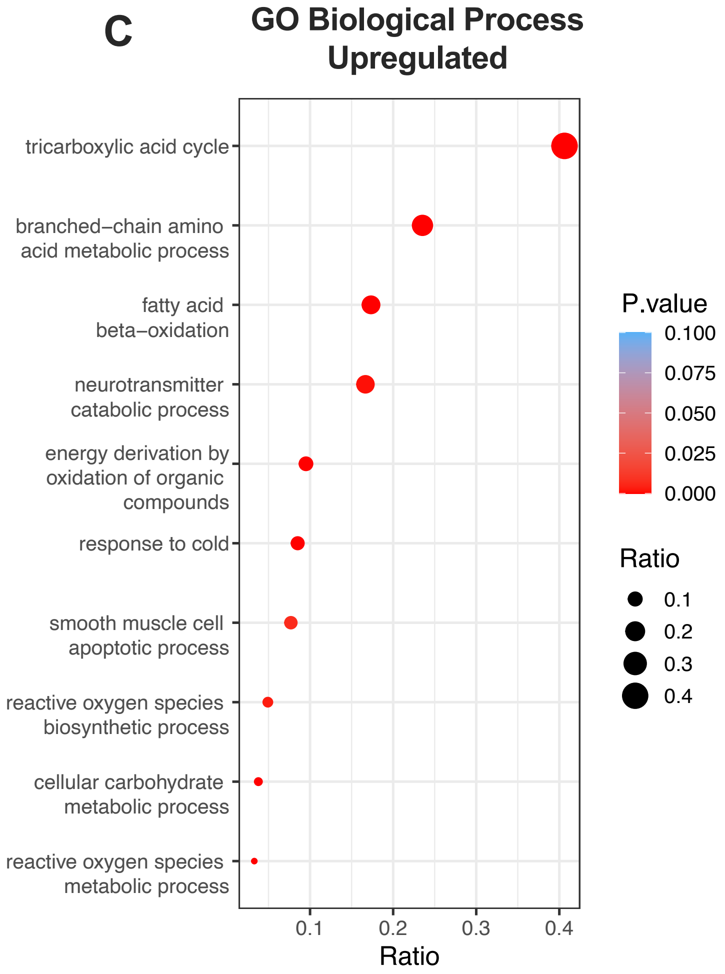
