## Supplementary figures and images for "Exercise Training and Cold Exposure Trigger Distinct Molecular Adaptations to Inguinal White Adipose Tissue"

### Supplemental Figure 2

#### Supplemental Figure 2

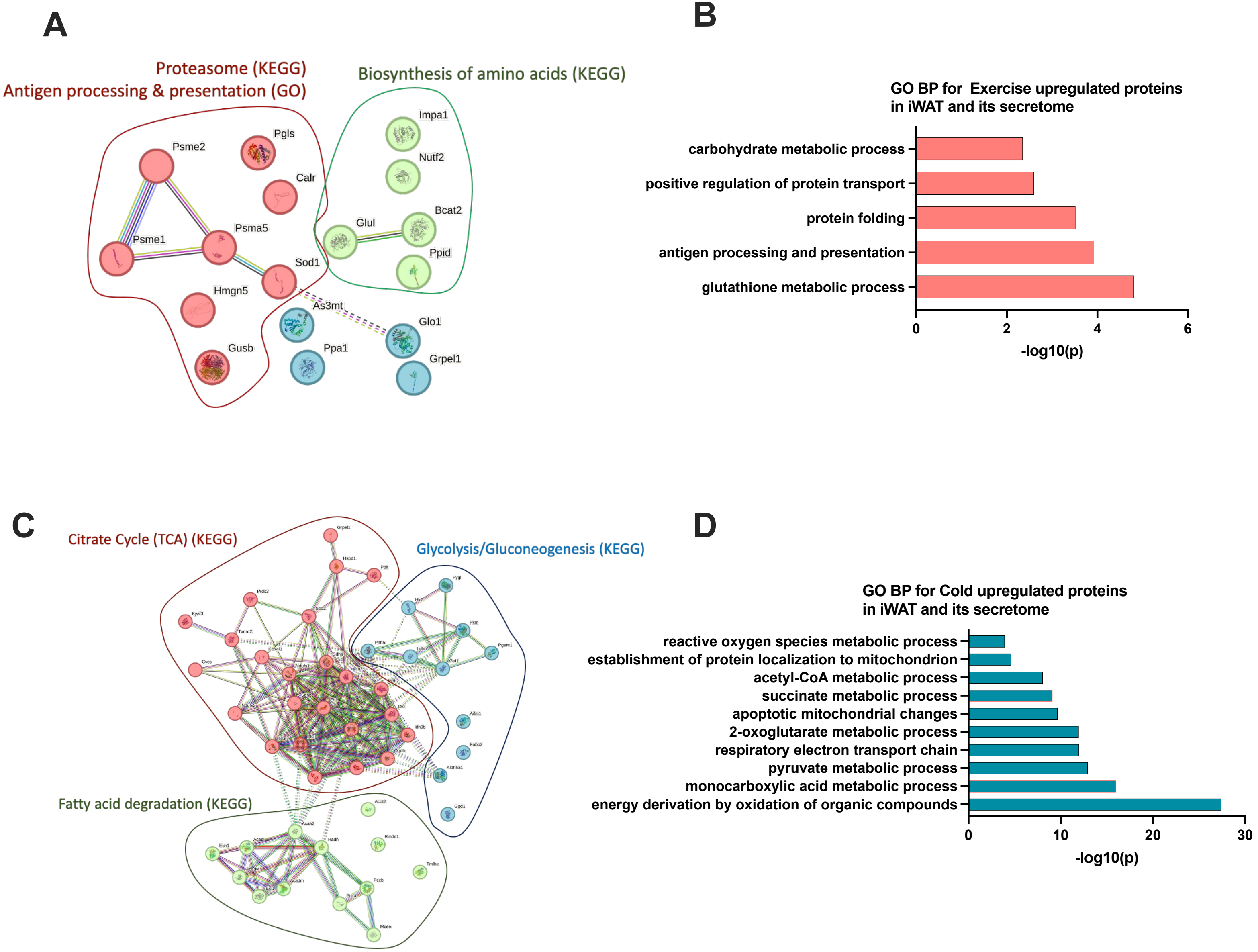

### Supplemental Figure 3

##### Supplemental Figure 3

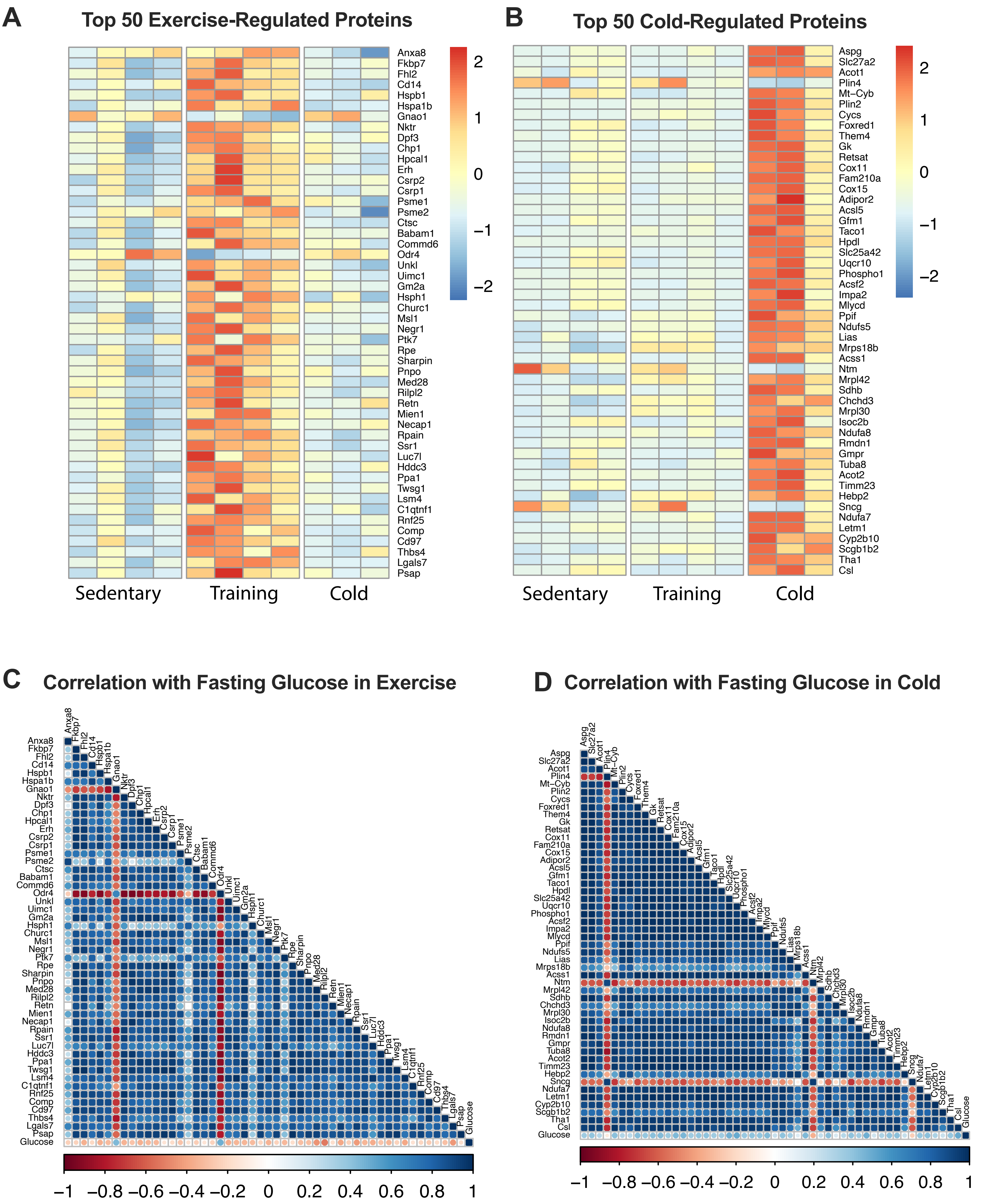
