## Supplemental Figure 4 for "Exercise Training and Cold Exposure Trigger Distinct Molecular Adaptations to Inguinal White Adipose Tissue"

**A**

| #Term ID | Term description | Strength | FDR |
|---|---|---|---|
| GO:0061951 | Establishment of protein localization to plasma membrane | 2.48 | 0.00036 |
| GO:0006904 | Vesicle docking involved in exocytosis | 2.41 | 0.0179 |
| GO:0009306 | Protein secretion | 2.13 | 0.0019 |
| GO:0006892 | post-Golgi vesicle-mediated transport | 2.07 | 0.0421 |
| GO:0007029 | Endoplasmic reticulum organization | 2.07 | 0.0421 |
| GO:0072659 | Protein localization to plasma membrane | 2.01 | 0.00015 |
| GO:0032869 | Cellular response to insulin stimulus | 1.98 | 0.0031 |
| GO:0006887 | Exocytosis | 1.81 | 0.0076 |
| GO:0017157 | Regulation of exocytosis | 1.79 | 0.0084 |
| GO:0048193 | Golgi vesicle transport | 1.78 | 0.0084 |
| GO:0006886 | Intracellular protein transport | 1.51 | 0.0019 |
| GO:0010256 | Endomembrane system organization | 1.51 | 0.0284 |
| GO:0051640 | Organelle localization | 1.48 | 0.0312 |
| GO:0000904 | Cell morphogenesis involved in differentiation | 1.42 | 0.0367 |

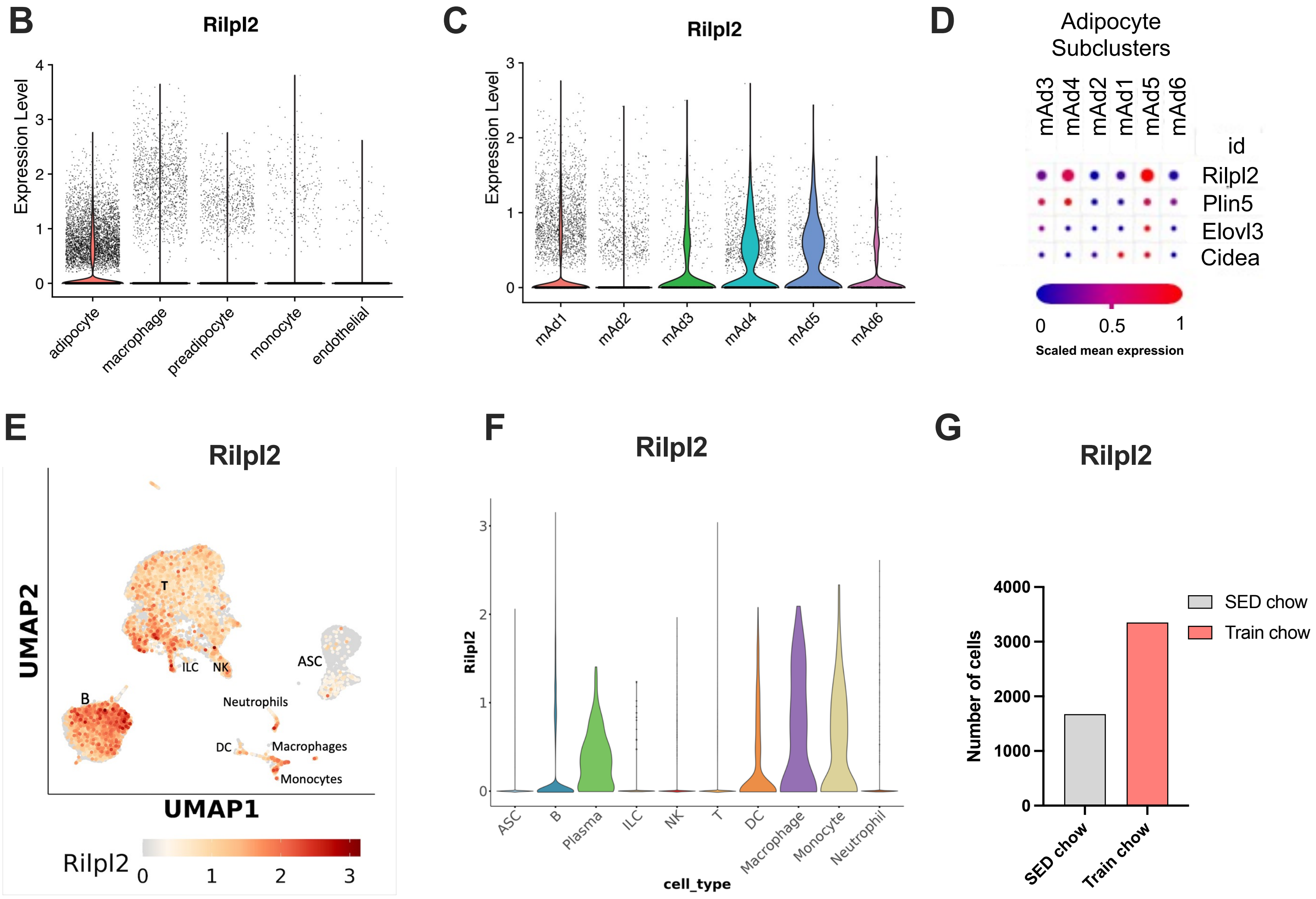
